# kinesin recruitment by adapter SKIP on melanosomes is dynamically controlled by LC3B phosphorylation

**DOI:** 10.1101/2021.03.11.434917

**Authors:** Yogaspoorthi Subramaniam, Divya Murthy, Desingu Ayyappa Raja, Amrita Ramkumar, Sridhar Sivasubbu, David G McEwan, Rajesh S Gokhale, Vivek T Natarajan

## Abstract

Anterograde melanosome transport is essential for adaptive skin tanning response. However, the molecular components involved, their interplay and regulation by external cues in melanosome transport remain under-explored. Silencing of kinesin motors revealed that several members including the established KIF5B and a novel candidate KIF1B, mediate melanosome movement. The camouflage behaviour of zebrafish embryos induced by incident light or α -MSH requires kif1b, suggesting a conserved melanosome transport machinery across vertebrates. Interestingly, the peri-nuclear melanosome accumulation upon kinesin knockdown is recapitulated by the silencing of autophagy effector MAP1LC3B (LC3B). Pull-down assays identified KIF1B, but not KIF5B, to be the LC3B-associated kinesin. LC3B binds the adapter SKIP via its LIR docking region that is proximal to Thr12 residue, a site for phosphorylation by Protein Kinase A. We demonstrate that phosphorylation of LC3B at Thr12 is stimulated by α-MSH, which potentiates the anterograde melanosome transport. Thereby, our study, identifies a novel kinesin motor KIF1B for melanosome movement and establishes LC3B as the key molecular component that facilitates α-MSH responsive mobilization of melanosomes.

**Key Highlights:**

- Kinesin screen reveals non-redundant use of KIF5B, KIF1B motors for melanosome transport
- kif1b is required for camouflage response in zebrafish and melanosome movement in mammals
- N-terminal region of LC3B interacts with adapter SKIP and couples kinesin KIF1B
- *α*-MSH activates PKA-mediated phosphorylation of LC3B to potentiate anterograde movement

**Significance:** Melanosomes are lysosome related organelles containing melanin pigment, that are synthesized in melanocytes and transferred to the recipient keratinocytes of skin. This involves long range melanosome movement within melanocytes to reach cell periphery for the transfer to follow. Physiologically, UV protection involves local secretion of melanocyte stimulating hormone (*α*-MSH) that acts on melanocytes to promote skin tanning response. Herein, we investigate the components involved in this process and establish that the melanosome movement is dynamically controlled by *α*-MSH through phosphorylation of LC3B. These findings establish the mechanism behind the rapid distribution of melanosomes during tanning response and provide opportunity to intervene for sun protection.

## Introduction

In human skin epidermis, incident solar radiation induces melanosomes to be rapidly transported from melanocytes to stratified keratinocytes. The transferred melanosomes aggregate around the nucleus to form a peri-nuclear cap protecting the genome from ultraviolet radiation-induced DNA damage. Rapid dispersion and aggregation of melanosomes in response to external cues is also well documented in early vertebrate melanocytes and serves the purpose of camouflage by adapting the animal to its immediate environment. Therefore, directed movement of melanosomes has been a subject of fascination for cell biologists and developmental biologists alike for this evolutionarily conserved process(Sköld, Aspengren et al. 2002, Barral and Seabra 2004, Hume and Seabra 2011). However, the identity of molecular components that mobilize the pigmented melanosomes, and its control by external stimuli have so far remained elusive.

The distribution of melanosomes is determined by the balance between anterograde movement by kinesins and retrograde movement by dynein. Kinesins tread melanosomes on the microtubule tracks present in the core of the cell, and the myosin dependent transport mediates movement on microfilaments at the cell periphery (Barral and Seabra 2004, Evans, Robinson et al. 2014, Oberhofer, Spieler et al. 2017). MYO5A motor moves melanosomes on peripheral microfilaments through RAB27A via the adaptor protein melanophilin (MLPH)(Wu, Bowers et al. 1998, Sckolnick, Krementsova et al. 2013). Centrality of each of these components in mobilizing melanosomes is evident in their mutant phenotypes that lead to depigmentation observed, both in patients with Gricelli’s syndrome as well as in animal models of the disease (Wilson, Yip et al. 2000, Matesic, Yip et al. 2001, Takagishi and Murata 2006).

Anterograde transport of organelles including melanosomes is mediated by the large superfamily of kinesin motor proteins (14 families). Kinesins are broadly categorised into 4 major groups based on their phylogenetic clustering: conventional kinesins, kinesin ll, Unc104/kinesin 1 and mitotic kinesins (Barral and Seabra 2004). Kinesin motors function in large complexes that confer selectivity of the cargo and cater to the cellular need to mobilize organelles. Study by (Ishida, Ohbayashi et al. 2015) elucidated the role of conventional kinesin, Kinesin 1 (KIF5B) along with the adapter protein PLKHM2/SKIP to facilitate anterograde movement of melanosomes, leading to the current understanding that this is the mediator of melanosome transport on microtubule tracks. Specific adapters such as FYCO1, JIP1 and SKIP facilitate cargo selection/ tracking by the kinesin (Kjos, Vestre et al. 2018). Thereby the identity of kinesins and associated adapters on melanosomes needs to be elucidated to decipher regulation of this controlled process.

In our previous work (Ramkumar, Murthy et al. 2017), we reported that the insertion of lipidated LC3B on melanosomal membrane is required for effective transport on microtubule tracks. In the current study we investigate members of the kinesin family and the associated adapter involved in LC3B-mediated melanosome transport. Six out of 38 kinesins were identified by our siRNA screen to alter the cellular localization of melanosomes. Biochemical pull-down studies of LC3B with kinesin candidates identifies KIF1B as the cognate motor. Involvement of kif1b in camouflage response of zebrafish embryos suggests conservation of the machinery involved in melanosome movement across vertebrates. Further, the role of SKIP as an LC3B-dependent adapter for KIF1B is delineated. Coupling of these components to *α*-MSH signalling pathway through LC3B phosphorylation, illustrates the external control of melanosome movement to the cell periphery in melanocytes. Hence our study establishes the role of LC3B, associated kinesin machinery and its regulation by melanocortin pathway to mediate long-range anterograde melanosome movement.

## Results

### Multiple kinesins are involved in the anterograde transport of melanosomes

Using an immunofluorescence-based co-localization study design, B16 mouse melanoma cells were examined for melanosome positioning on tubulin tracks following knock-down of specific kinesin members in a targeted kinesin screen. A custom-made siRNA library against kinesins was designed and 40 out of 44 kinesins in the mouse genome could be individually targeted (**Table S1**). Subsequently, the cells were stained for microtubule tracks using tubulin antibody, for melanosomes using HMB45 antibody and were subsequently evaluated melanosome positioning within the cell. High content microscopy was employed to capture and analyse around 100 cells per treatment. Silencing of two of the kinesins *Kif21b* and *Kif3b* resulted in cell death, and the melanosome positioning could not be evaluated. The first channel set to detect tubulin was taken for autofocus, and the second channel was set for melanosomes stained with HMB45. HMB45 was chosen as it robustly marks the melanosomes from stage I to late stages of maturation (Taatjes, Arendash-Durand et al. 1993). The positioning of melanosomes was analysed by tracing two concentric circles across the cell and the area normalized intensity ratio of HMB45 staining between the inner circle and the outer ring was calculated (detailed in materials and methods section) (Fig 1A). While the control cells had ratios closer to 1 suggesting an overall uniform distribution, greater ratio in several kinesin silenced conditions (<1 or <0.5) indicated peri-nuclear aggregation of melanosomes. Following this analysis, we found that silencing 6 out of 38 screened kinesins results in peri-nuclear positioning of melanosomes (Fig S1A & S1B). These are – *Kif1a, Kif1b, Kif2b, Kif5b, Kifc1* and *Kifc3*. While the two mitotic kinesins *Kif11* and *Kif24*, did not affect melanosome positioning, *Kif2b* speculated to be a mitotic centromere associated kinesin, seemed to have a significant effect (Miki, Setou et al. 2001). Interestingly, we observed higher colocalization between melanosomes and tubulin in most of the conditions wherein the ratios were greater than 1, suggestive of melanosome accumulation on tubulin tracks. Hence, based on this kinesin screen we establish the previously demonstrated kinesin motor KIF5B to be one of the kinesin motors for melanosomes transport (Ishida, Ohbayashi et al. 2015). Further, this screen identifies other kinesin family members that could tread melanosomes on tubulin tracks and suggests that they do so in a non-redundant manner as silencing of single member individually results in the aggregation phenotype.

**Figure 1:**
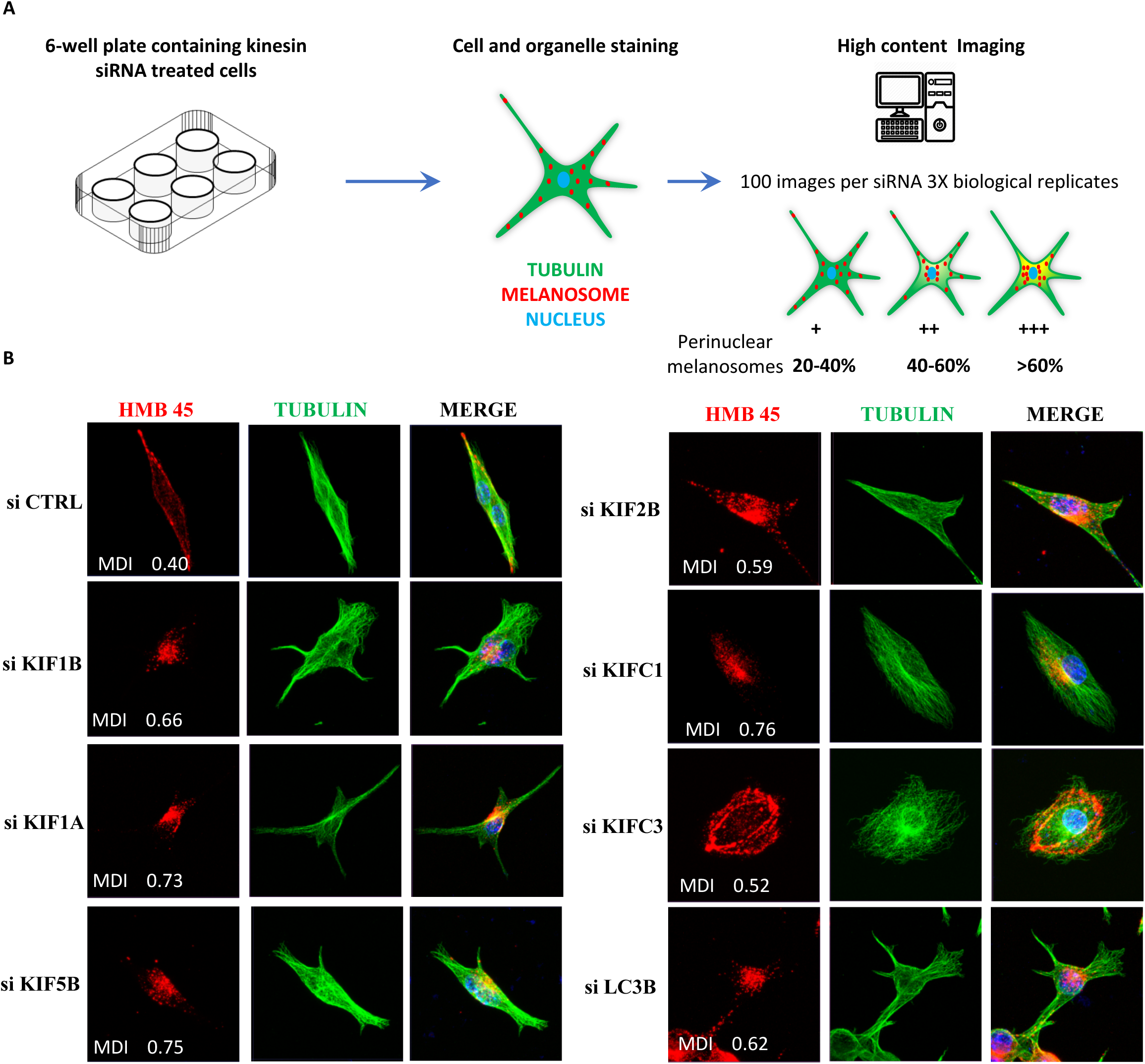
Kinesins involved in anterograde melanosome transport. A. Schematic of experimental set-up. Cells are immune-stained with HMB45 –(Red) marking melanosomes and Tubulin-(Green) marking microtubule tracks. 100 images were analysed per siRNA. The ratio of HMB45 staining in the centre of the cell (Ci) to the intensity in the ring region (Ri) normalized to the respective area. n=3. B. Representative images of melanosome positioning upon siRNA-mediated silencing of individual kinesins.

Recently, the C-terminal region of KIFC1 was shown to specifically localize to melanosomes in melanocytes and melanocores, the melanosomes transferred from melanocytes to keratinocytes (Ishida, Marubashi et al. 2017). In our study, silencing of *Kifc1* resulted in partial peri-nuclear accumulation in melanocytes, which is in agreement with KIFC1 and KIFC3 involvement in retrograde transport (Hirokawa, Noda et al. 2009). KIF2B is a lesser known kinesin involved in microtubule remodelling and chromosome segregation (Kline-Smith and Walczak 2004). Interestingly, we found two of kinesin-3 motors, KIF1A and KIF1B, to feature among the hits. KIF1A, the most studied member of this family, carries a sub-set of synaptic vesicles containing synaptogamin, synaptophysin and Rab3, across the neuronal axons (Okada, Yamazaki et al. 1995). KIF1B is a related kinesin and is believed to function like KIF1A and is associated with the transport of mitochondria and lysosomes (Guardia, Farias et al. 2016).

To validate the screen and rule out changes in microtubule tracks as a reason for melanosome aggregation, we performed high magnification confocal microscopic evaluation of the silenced cells from promising hits (Fig 1B). Upon silencing of *Kif1a, Kif1b, Kif2b, Kif5b, Kifc1* and *Kifc3* with a sequence independent SMART pool siRNA (Dharmacon) targeting each of them individually, there were no apparent disruptions in microtubule tracks and the perinuclear aggregation was evident. Thereby these kinesins are involved in altering the position of melanosomes. While the peri-nuclear melanosome aggregation phenotype was stark in all the hits, it was less evident in the KIF2B silenced cells. Silencing of *Kifc3* resulted in the peri-nuclear accumulation of melanosomes distinct from other conditions, where the distribution was clearly ring-like. This suggested a possible arrest in the transport from actin-microtubule junctions in the periphery of the cell due to a lack of retrograde transport and recapitulates the phenotype of *Atg4b* silencing reported earlier (Ramkumar, Murthy et al. 2017). Given their role as retrograde motors, perinuclear melanosome accumulation upon *Kifc1* silencing is surprising. A similar perinuclear accumulation of peroxisomes could be rationalized by a complex interplay involving retrograde kinesins with KIF5B and Dynein (Dietrich, Seiler et al. 2013, Guardia, De Pace et al. 2019). Hence the major kinesins involved in the anterograde transport are KIF1A, KIF1B, KIF5B and KIFC1. Among these four kinesins KIF1A, KIF1B and KIF5B are known to mobilize lysosomes which closely resemble melanosomes due to their common origin and shared proteome (Vickrey, Bruders et al. 2018). Hence, we focused further on the role of KIF1A and KIF1B on melanosome dispersion further in a distinct model system.

### Kif1a/Kif1b mediate long-range dispersion of melanosomes during zebrafish camouflage behaviour

Zebrafish embryos respond to external light cue and reorganize their melanosomes according to illumination of their immediate environment. This adaptive camouflage response is mediated by the coordinated aggregation or dispersion of melanosomes on microtubules. This behavioural phenotype permits the study of coordinated long range melanosome movement and to investigate the associated machinery (Rodionov, Hope et al. 1998, Rogers and Gelfand 1998).Orthologs of LC3B as well as kinesins kif1a and Kif1b are well conserved in zebrafish (Fig S2A). We evaluated the ability of zebrafish larva to mediate melanosome movement upon knockdown of kinesins *kif1b* and *kif1a* (Fig 2 A&B). Zebrafish embryos kept under dark background conditions result in melanosomal dispersion and the melanophore (melanocyte) appears spread out, this phenomenon can be recapitulated by treating the embryo with the melanocortin activator α-MSH (Fig 2C).

**Figure 2:**
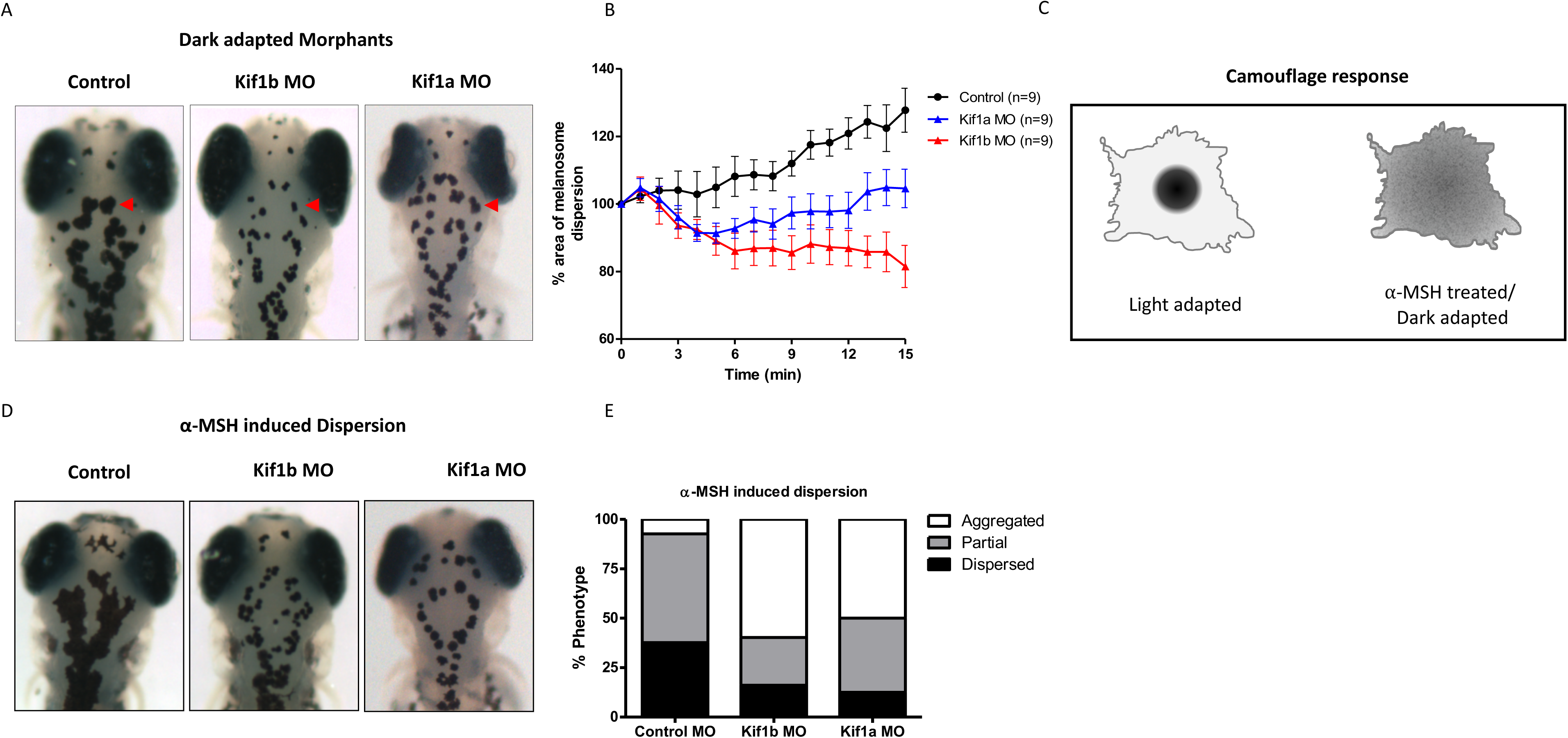
Kinesins Kif1a and kif1b mediate melanosome dispersion in Zebrafish camouflage response. A. Dorsal head of the morphant and control zebrafish embryos at 4dpf after dark adaptation. the pigmented black cells here are the melanophores. B. Live imaging of morphant and control zebrafish embryos was performed after light adaptation in a minimally illuminated conditions and the anterograde melanosome dispersion was captured as percentage are of melanosome dispersion within the melanophores (15 melanophores per animal and 9 animals per group). The graph depicts the percentage area of melanosome dispersion upon dark adaptation. Bars represent mean±SEM. Control vs Kif1a MO = ** (p Value=0.011); Control vs Kif1b MO = **** (<0.0001) C. Diagrammatic representation of melanosome dispersion during camouflage behaviour in response to physiological cues such as alpha-MSH and light/dark adaptation. D. Representative images of dorsal head regions of 4dpf zebrafish embryos treated with 15uM of α-MSH to induce dispersion of melanosomes in melanophores. E. Quantification of the α-MSH induced dispersion phenotype. The embryos were categorized as dispersed, partial or aggregated. The embryos were categorized as Dispersed partial and aggregated based on mean Gray values quantitated from 8-bit images of 4% PFA fixed embryos. The region between eyes of the embryo till the otic region was selected for analysis. Mean grey values of 40-60 was categorized as Dispersed, 60-80 as Partial and 80-100 as Aggregated. At least 20 embryos were taken for each morphant (n=2).

We performed assays on control, *kif1a* and *kif1b* morphants (MO) zebrafish embryos at 4 days post fertilization (dpf). At this stage of development, the embryos display robust background adaptation response. We utilized a surrogate phenotype to score *kif1b* morphants and segregated embryos based on touch stimulus based response (Lyons, Naylor et al. 2009). The embryos were dark adapted, paraformaldehyde fixed, and bright field imaging was carried out. While the melanosomes in control embryos are well dispersed, both the *kif1a* and *kif1b* morphants showed a distinctive aggregation phenotype and the melanosomes appear to be concentrated (Fig 2A). Since in the control animals the response is quick and adaptation is complete by around 10 – 15 minutes, we developed a kinetic assay to quantitate the dispersion of melanosomes.

In the live imaging set up zebrafish embryos were light adapted, which led to the melanosomes being aggregated. Subsequently, the immobilized embryos were imaged in bright field, under limited light exposure. Kif13a, a kinesin known to affect melanosome biogenesis but not the melanosome positioning based on our B16 kinesin screen, was included as a control in this study (Delevoye, Hurbain et al. 2009). *kif13a* morphants showed a noticeable reduction in the melanin content per melanophore but had sufficient melanin content in them for imaging. We observed that the control as well as *kif13a* morphants showed a 30% increase in area occupied by the melanosomes by around 15 min (Fig S2B). However, in both *kif1a* and *kif1b* morphants the area covered by melanosomes did not increase and rather showed a downward trend. In the *kif1a* morphants the melanosomes return to the original position. In the *kif1b* morphants we observe that the area covered by melanosomes shows a significant decrease of around 20% (Fig 2B) (**Videos 1 and 2**).

The camouflage behaviour is a response controlled by the intracellular cAMP levels. Melanosome dispersion is primarily governed by melanocyte stimulating hormone, α-MSH. Therefore, we treated the light adapted embryos with α-MSH in dark for 10 min and fixed them before imaging. Upon exposure to α-MSH, the control animals demonstrated a complete dispersion of melanosomes whereas the *kif1a* and *kif1b* morphants were unable to disperse the melanosomes and they remained aggregated (Fig 2D&E). These observations thus reveal the importance of both the kinesins identified by siRNA screen. Kif1b as well as Kif1a are necessary for long range melanosome movement in melanocytes and are responsible for the zebrafish camouflage behaviour.

### Kinesin KIF1B physically interacts with LC3B

Striking similarity between the kinesin and *Lc3b* silencing on the positioning of melanosomes suggested that the two machineries could possibly cooperate for the anterograde melanosome movement. Tethering of motors on cargo vesicles is mediated by several modes of interaction. In the context of melanosomes, melanophilin present on melanosome surface connects with the membrane through lipidated RAB27A and interacts with myosin Va to facilitate the transport on microfilaments. KIF5B complexes with lipidated RAB1A present on melanosomes for movement on microtubules (Ishida, Ohbayashi et al. 2012). In these lines, LC3B by virtue of being lipid anchored offers a key role as a possible tether. Our observation that silencing of LC3B phenocopies that of kinesins, along with its ability to anchor complexes, makes LC3B a plausible candidate for binding the kinesin motor.

To identify kinesin interacting partners of LC3B we performed pull-down studies with lysates of melanosomes isolated from pigmented B16 melanoma cells. Either recombinant GST alone or LC3B-GST fusion protein was immobilized on glutathione beads and incubated with melanosome lysate and the specifically bound proteins were eluted with reduced glutathione. Proteomic analysis revealed a set of proteins to be bound to GST-LC3B (**Table S2**) Prominent among them are the tubulin subunits, which is not surprising as LC3B was initially identified as a tubulin interacting protein and is appropriately named as microtubule associated protein 1 light chain 3B (MAP1LC3B). Some of the cofractionating kinesins KIF1A, KIF1B KIF7, KIFC3 also featured albeit with low coverage in the pull-down list with at least one peptide picked up in the mass spectrometric detection (**Table S2**). Since KIF7 did not show melanosome aggregation phenotype and KIFC3 showed a ring-like phenotype, we concentrated on KIF1A and KIF1B. As the beta isoform of KIF1B is almost identical to KIF1A, they are likely to function similarly in treading lysosomes. Additionally, in an earlier study we had indications of KIF1B to be coregulated with pigmentation and hence this kinesin emerged as a strong candidate (Raja, Gotherwal et al. 2020).

### LIR mediated interaction by LC3B mediates melanosome positioning

To investigate the interaction of kinesin with LC3b, we performed western blot analysis on the pull-down with KIF1B antibody and for comparison analysed KIF5B, the established melanosome mobilizing kinesin. While KIF1B specifically bound GST-LC3B but not GST, KIF5B could not be detected in the pulldown, suggesting that KIF1B is the cognate binding kinesin of LC3B (Fig 3A). Effectors typically associate with LC3B using the LIR docking site (LDS) on LC3B (Fig 3B). Using site-directed mutagenesis we generated the mutant form of LC3B which lacks the regions necessary for LIR interactions by deleting the first 10 residues, LC3BΔN10 (Shvets, Fass et al. 2008). We then investigated the ability of siRNA-resistant wild type *LC3B* (human ortholog) or this mutant form to rescue the melanosome aggregation phenotype observed upon *Lc3b* silencing. While the wild type LC3B could restore melanosome distribution as demonstrated previously in our earlier study (Ramkumar, Murthy et al. 2017), the LC3BΔN10 mutant form could not distribute the melanosomes and a peri-nuclear accumulation could still be observed in a substantial number of cells (Fig 3C & 3D). These findings substantiate the ability of LC3B to interact with associated proteins using its N-terminal LIR docking site (LDS) and thus, the interaction is critical to mediate the effects on melanosomal positioning in melanocytes. Therefore, we speculate that a LIR dependent interaction is essential to mediate movement of melanosomes. Given the other interactions of LC3B in vesicular dynamics, it is likely that the interaction is indirect and mediated by an adapter protein.

**Figure 3:**
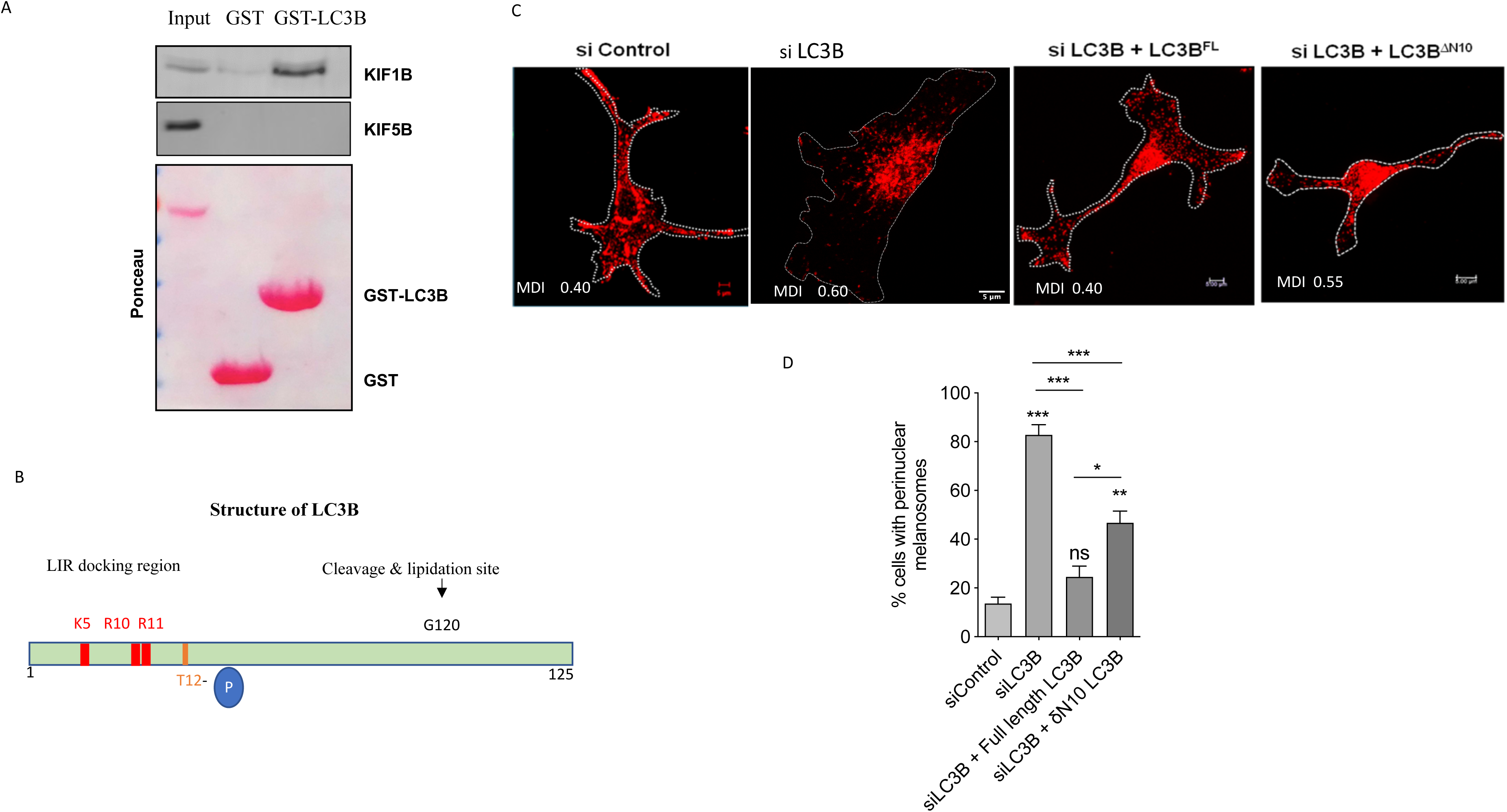
LC3B binds to KIF1B motor and LIR-Docking site in LC3B is required to facilitate the anterograde melanosome dispersion. A. Recombinant GST-LC3B interacted with melanosome lysate was subjected to western-blot analysis and probed with Kif1b and Kif5b antibodies. Ponceau S staining of the membrane was used to normalize the pull-down efficiency. B. Schematic representing LC3B domain architecture depicts the starting N-terminal 12 amino -acids containing LIR docking site (LDS), the proximal site for PKA mediated phosphorylation Thr12 and the C-terminal cleavage and lipidation site at Gly120. C. Melanosome distribution based on HMB45 (Red) immuno-staining in endogenous LC3B silenced cells complemented with either siRNA-resistant LC3B (full length) or LC3B*Δ*N10. Scale bar −5uM. D. Bar graph representing the quantitation of percentage analysed cells that retain peri-nuclear melanosome aggregation. (n<50 cells) Bars represent mean ±s.e.m. across replicates.

### SKIP is a common adapter for both Kif5b and Kif1b

Kinesins are recruited through adapters such as APP (Kamal, Stokin et al. 2000), FYCO (Pankiv, Alemu et al. 2010), JIP1 (Verhey, Meyer et al. 2001) and SLP1 (Arimura, Kimura et al. 2009) either directly or as a part of a complex for mediating organelle movement. Interestingly, lipidated LC3B is known to be present in such complexes and hence could have a functional role in mediating the movement of organelles (Fu, Nirschl et al. 2014, Sakurai, Tomita et al. 2017). LC3B in complex with FYCO1 interacts with kinesin(s) and facilitates plus-end directed movement of autophagic vesicles (Pankiv, Alemu et al. 2010).

To identify the possible adapter for KIF1B on melanosomes, we first interrogated SKIP, the adapter that connects KIF5B to RAB1A. Towards this, we first overexpressed HA-tagged SKIP in B16 melanoma cells and examined its colocalization with the melanosome marker, HMB45. Considerable colocalization of SKIP and HMB45 at the dendritic tips of melanocytes reaffirmed the previous observations by (Ishida, Ohbayashi et al. 2015)(Fig 4A, Fig S3C). The role of SKIP as a melanosome adapter is further corroborated by perinuclear aggregation of melanosomes upon SKIP down regulation, a phenotype strikingly similar to LC3B and kinesins KIF5B/1A/1B knockdown (Fig 4B & C).

**Figure 4:**
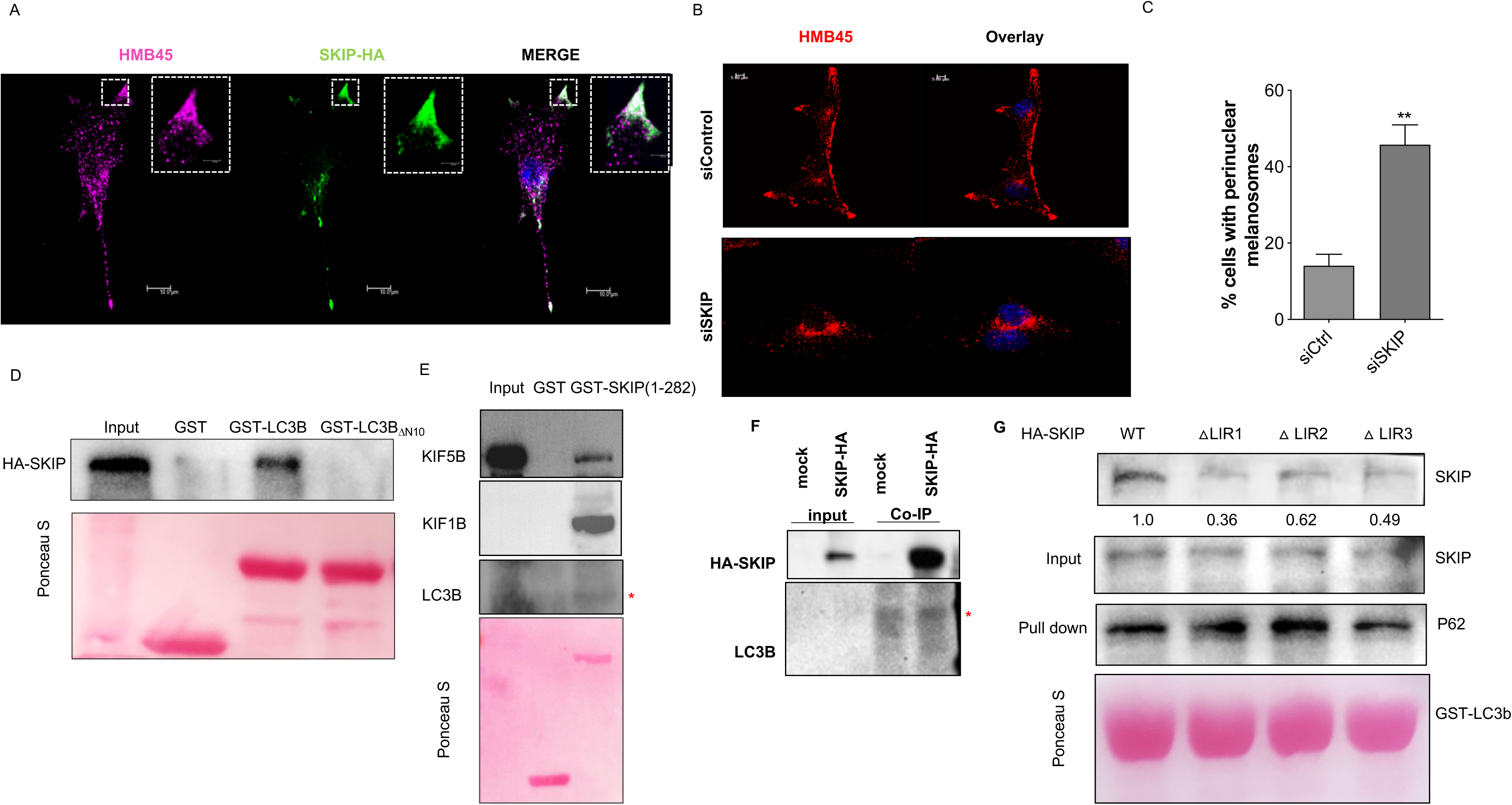
LC3B via adaptor protein SKIP recruits Kif1b to melanosomes. A. Colocalization analysis of B16 cells transfected with HA-tagged SKIP and immune-stained with anti-HA antibody (Purple) and HMB45 antibody (Green) marking melanosomes. Scale bar-10uM. B. Melanosome distribution based on HMB45 (Red) immuno-staining and DAPI marking the nucleus upon SKIP silencing. Scale bar-5uM C. Bar graph representing the quantitation of cells exhibiting peri-nuclear melanosome aggregation phenotype. D. Recombinant GST-LC3B, GST-*Δ*N10-LC3B or GST were incubated with lysate from Human embryonic Kidney (HEK) cells expressing HA-SKIP. Western blot analysis after pull-down was probed for SKIP using anti-HA antibody. E. GST-SKIP or GST interacted with melanosomal lysate and the pulled-down was probed with anti-Kif1b antibody, anti-Kif5b antibody and anti-LC3B antibody. Ponceau staining is used for normalisation. Red asterix indicates a non-specific band of IgG light chain. F. Lysate from HEK293T expressing HA-SKIP were immunoprecipitated with anti-HA agarose beads and probed for LC3B by western-blotting. G. GST-LC3B or GST-empty vector interacted with LIR motif mutants of SKIP. Western blot analysis after pull-down was probed for SKIP using anti-HA antibody, and for p62. Ponceau staining is used for normalisation.

To examine if SKIP acts as the adaptor for LC3B and KIF1B interaction, we first performed GST-LC3B pulldown assay with cell lysate from HA-tagged SKIP-expressing cells. We observed interaction of GST-LC3B with SKIP as well as P62 while these proteins were not associated with GST (Fig 4D). Further to confirm this interaction we performed pull down using GST-SKIP (RUN domain) and observed that it interacts with both the kinesins KIF1B and KIF5B (Fig 4E). Interestingly, the enrichment of KIF1B is several folds higher suggesting a stronger interaction. A faint but consistent interaction with LC3B could also be detected, which was strengthened by the coimmunoprecipitation of SKIP with LC3B (Fig 4F). As the N-terminal deletion mutant of *LC3B* was unable to rescue melanosome dispersion, we speculated that this region is involved in complexation (Fig 3C). The cognate mutation as a GST fusion protein failed to effectively pull-down HA-SKIP (Fig 4F). Corroborating this, different LIR motif mutants of SKIP failed to interact with GST-LC3B (Fig 4G), further substantiating that SKIP-LC3B interaction is indeed mediated by LIR domains on SKIP. SKIP LIR1 (Fig S3B) is the dominant LIR motif that accounts for SKIP interaction with LC3B LDS.

These results indicate the existence of a melanosome transport machinery involving LC3B as the melanosomal anchor, SKIP as the intermediate adaptor and KIF1B as the motor protein that treads melanosomes on microtubules. Interestingly, the same adapter SKIP is involved with both the kinesins KIF5B as well as KIF1B that mediate melanosomal transport, however the anchorages are seemingly distinct. This shared adapter may be a mechanism to facilitate rapid switching between kinesins that preferentially tread on distinct subtypes of microtubule tracks (Guardia, Farias et al. 2016).

### α-MSH treatment increases the speed of melanosome movement in mammalian melanocytes

While the camouflage behaviour of zebrafish is well documented and the role of MSH is unequivocal, whether a similar response is observed in mammalian melanocytes is not clear. In earlier studies, authors used murine melanocytes alone and in coculture with keratinocytes to evaluate melanosome transfer (Virador, Muller et al. 2002, Ma, Ma et al. 2014). It was observed that α-MSH stimulated the release of melanosomes to the extracellular milieu and concomitantly elevated several genes involved in the process (Virador, Muller et al. 2002). However, the status of cellular changes on melanosome motility has not been systematically studied in response to MSH. Hence, we resorted to study melanosome movement in B16 melanoma-derived cells by live imaging. B16 cells transfected with OA1-GFP which has been shown previously to mark melanosomes was employed in this study (Bruder, Pfeiffer et al. 2012).

Corroborating the screen and subsequent high magnification imaging, both kinesins as well as *Lc3b* silencing [as reported in our previous study (Ramkumar, Murthy et al. 2017)] resulted in massive perinuclear aggregation of OA1-GFP positive melanosomes (**Video 3-6**). There are fewer melanosomes distributed in the rest of the cell. The frequency of bidirectional movement among these isolated melanosomes appears to be low and majority of them showed retrograde movement. As the silencing resulted in dense perinuclear aggregation, neither the directionality nor the speed of movement could be calculated from the videos. These observations were common to both the kinesins as well as *Lc3b*, establishing that the perinuclear aggregation is due to compromised anterograde movement and also indicated that the two machinery may physically or functionally interact to mobilize melanosomes. We then imaged B16 cells for 5 minutes as control and for α-MSH treatment, we treated cells with 10μM α-MSH and then imaged for 5 minutes. In successive frames melanosomes were tracked and the speed of movement was quantified. We observed that stimulating the cells with α-MSH potentiates the speed of moving melanosomes (Fig 5A).

**Figure 5:**
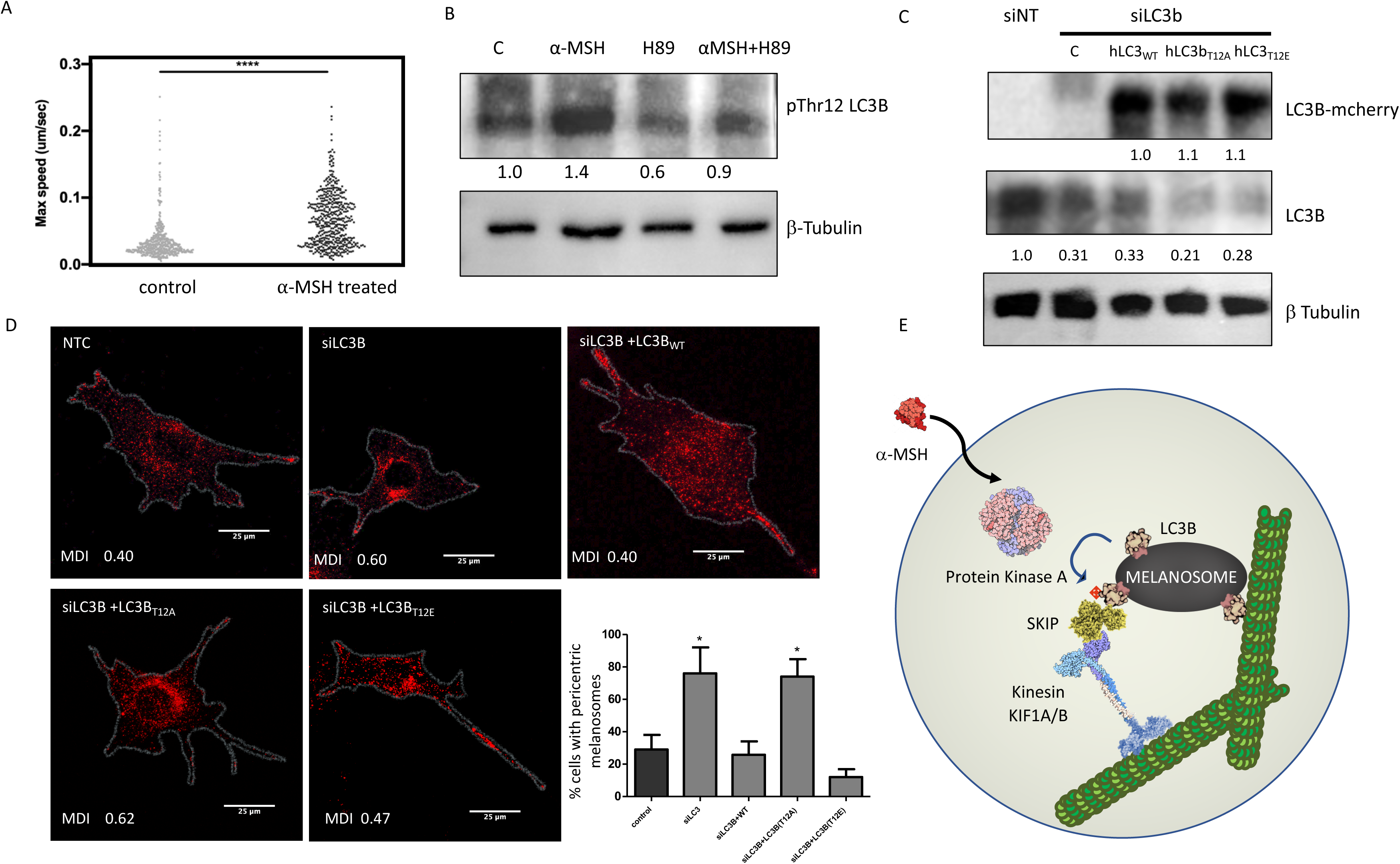
PKA mediated phosphorylation of LC3B is essential for LC3B facilitated melanosome transport. A. Speed of melanosome movement calculated from time-lapse imaging of melanosomes tagged with OA1-GFP upon 10µM *α*-MSH treatment. 200 melanosomes per cell were tracked and 3 cells per treatment condition were imaged. Bar represents the mean±s.e.m. **** depicts a p-value <0.0001. Wilcoxon test was applied. B. B16 cells were treated with 1µM *α*-MSH or 1µM H89 or the combination of both. Lysate prepared from these cells were analyzed for phosphorylation of LC3B at Thr12 using anti-PhosphoLC3B-Thr12 antibody and normalized to beta-tubulin. n=3. Tukey’s multiple comparisons test was applied and p-value <0.05 is considered significant. C. Site directed mutagenesis was used to generate hLC3B^T12A^ (non-phosphorylatable) and hLC3B^T12E^ (phospho-mimetic). B16 cells were co-transfected with siLC3B and siRNA resistant human mCherry-tagged LC3B or hLC3B^T12A^ or hLC3B^T12E^. LC3B silencing by siRNA and rescue by expression of respective resistant human constructs were analyzed using western blot with anti-LC3B antibody and mCherry antibody. Fold change was calculated with respect to beta-tubulin. n=3. Tukey’s multiple comparisons test was applied and p-value <0.05 is considered significant. D. Melanosome distribution based on HMB45 immuno-staining upon silencing of endogenous LC3B and complementation with either siRNA resistant human mCherry tagged LC3B or hLC3B^T12A^ or hLC3B^T12E^. MDI=melanosome dispersion index. Scale bar=25uM. n=30 cells. Tukey’s multiple comparisons test was applied and p-value <0.05 is considered significant. E. Bar graph depicts the percentage of cells with peri-nuclear melanosome aggregation in siLC3B cells rescued with either full length, phosphomimetic or non-phosphorylatable human LC3B constructs. n=30 cells. Bars represent mean±s.e.m. Tukey’s multiple comparisons test was applied and p-value <0.05 is considered significant. F. Schematic model depicting the components – melanosome tethered LC3B, kinesin motor KIF1B and the adaptor SKIP, involved in anterograde melanosome movement. Dynamic regulation of this movement is mediated by MC1R activation. PKA activated by MC1R mediates phosphorylation of LC3B that facilitates anterograde melanosome movement by effective engagement of SKIP and recruitment of KIF1B. Efficient assembly of stable transport machinery is important for long-range anterograde melanosome movement in response to physiological cues.

### α-MSH mediates LC3B phosphorylation via PKA and potentiates the melanosome movement

Having identified the components, LC3B, the adapter SKIP and the kinesin KIF1B involved in the melanosome movement, we then resorted to investigate the mechanism by which α-MSH potentiates the speed of movement by modulating this machinery. Addition of α-MSH results in a rapid mobilization of melanosomes in B16 cells, necessitating identification of the underlying mechanism behind this rapid response in melanocytes (Fig 5A). Pro-pigmenting effect of α-MSH is well documented (Praetorius, Grill et al. 2013, Mujahid, Liang et al. 2017), but these are mediated by the transcriptional response elicited through the central transcription factor MITF. However, we observed that α-MSH treatment increased melanosome speed (Fig S4), but it was rapidly reversed to basal levels after around half an hour of treatment, hence we focussed on pathways upstream of MITF that could catalyse this rapid response.

LC3 family proteins are known to be phosphorylated at Ser/Thr residues by mTOR responsive STK4 kinase as well as Protein Kinase A (PKA) at two distinct sites (Cherra, Kulich et al. 2010, Wilkinson, Jariwala et al. 2015). α-MSH activates MC1R which activates PKA by increasing intracellular cAMP levels, rendering this as a possible link. *In-vitro* phosphorylation reaction was set up with GST-LC3B as well as the GST-LC3B LDS mutant ΔN10. Upon probing with Thr12 phosphorylated LC3B antibody, increased phosphorylation was observed for the full-length protein by PKA (Fig S5). Interestingly, the Thr12 residue is close to the LDS and hence alterations in mutant ΔN10 could deter phosphorylation. We then went on to decipher whether this phosphorylation could be achieved by α-MSH-mediated activation of melanocytes. B16 cells were treated with α-MSH for 30 min or pre-treated for 2h with H89 a selective inhibitor of PKA or as a combination. MSH treatment increased phosphorylation of LC3B and the basal as well α-MSH stimulated phosphorylation was suppressed by H89 (Fig 5B). Evidently by this mechanism α-MSH could stimulate speed of melanosome movement.

To test whether phosphorylation has an effect on melanosome mobilization, we resorted to the rescue experiment with either non-phosphorylatable or the phosphomimetic mutant of LC3B reported earlier (Cherra, Kulich et al. 2010). Silencing of endogenous LC3B using siRNA resulted in a decrease of around 75% (Fig 5C). Overexpression of mCherry-tagged siRNA-resistant human *LC3B* was capable of rescuing the melanosome aggregation phenotype. The non-phosphorylatable T to A (LC3B_T12A_) mutant was unable to rescue the phenotype whereas phosphomimetic T to E (LC3B_T12E_) mutant was capable of mobilizing the melanosomes comparable to the wild type (Fig 5D). As the LDS is close to the site of phosphorylation it is likely that the complexes are differentially stabilized by the event of phosphorylation and result in facilitated anterograde movement. Thereby this study provides a mechanistic basis to α-MSH-mediated melanosome movement facilitated by the LC3B, kinesin KIF1B and adaptor SKIP complex.

## Discussion

Melanosomes offer a paradigm to understand the transport of organelles within and also across cells. While the dynamic movement of melanosomes is well studied, it is rather surprising that the identity of melanosome treading machinery has remained enigmatic. Based on nocodazole mediated disruption of microtubules, studies demonstrate that actin tracks enable anterograde melanosome movement (Aspengren, Wielbass et al. 2006). This has led to the notion that MYO-VA is the key anterograde motor. However, melanosomes in the dilute mutant mouse (MYO-VA mutant) melanocytes do demonstrate anterograde movement indicating the role for other anterograde motors such as kinesins (Hume, Ushakov et al. 2007). Studies in late 1990s indicated the role of kinesin II motor in melanosome transport (Tuma, Zill et al. 1998, Rogers, Karcher et al. 1999). This primarily involved microinjection of antibody against kinesin II to block the movement of melanosomes. Later,(Hara, Yaar et al. 2000) demonstrated the role of Kinesin 1 in long range melanosome transport.

It is surprising that several kinesins when individually knocked down in melanocytes give rise to perinuclear melanosome aggregation. We speculate that the anterograde movement of melanosomes on microtubule track is a cooperative interplay of at least two different families, kinesin I (KIF5B) and Kinesin III (KIF1B and KIF1A). Mechanistically this model is supported by preferential movement of lysosomes on acetylated tracks by KIF1A/B and on tyrosinated tracks by KIF5B (Guardia, Farias et al. 2016). SKIP emerged as a common adapter to both KIF5B as well as KIF1B, but with distinct anchorages to melanosomes, RAB1A and LC3B respectively. While KIF1B can directly bind lipids with its Plekstrin Homology (PH) domain, KIF5B is assisted by the PH domain present in the adapter protein SKIP (Guardia, Farias et al. 2016). This shared adapter may recruit distinct kinesins and permit easy switching of tubulin tracks during movement. Therefore, the kinesin motors are likely to cooperatively move melanosomes. Hence, we propose a model wherein the adapter SKIP acts as a fulcrum for alternating between the two kinesin motors and provide cooperativity for the movement of melanosomes (Fig 5E).

The most exciting observation in this study emerged as the phosphorylation of LC3B by the α-MSH pathway. This provides a mechanistic basis of dynamic coupling of melanosome movement to external pro-pigmenting cues. It is interesting to note that α-MSH secretion is induced by the exposure of skin to UV rays. α-MSH mediates the induction of pigmentation genes by the central melanocyte transcription factor MITF to facilitate fresh synthesis of pigmented melanosomes (Kawakami and Fisher 2017). This would entail a lag between initial UV exposure and the tanning response. Coupling of LC3B phosphorylation to MSH pathway provides the opportunity to mobilize existing melanosomes and provide immediate protection, which could then be backed up by fresh melanin synthesis.

Thereby, in this study we identify several components of the machinery that mobilize melanosomes on microtubule tracks. Further, we establish the physical interaction between the LC3B on melanosome membrane with the kinesin KIF1B on tubulin track via the adapter SKIP. Dynamic regulation of this complex mediated movement via phosphorylation of LC3B provides a direct mechanism to control the anterograde melanosome transport. Thus far dynamic mobilization of autophagic vesicles is known only in conditions such as nutrient starvation. Herein utilizing shared components of the autophagy pathway and coupling it to the melanocortin signalling, pigmentary system seems to have evolved a rapid melanosome dispersion modality for photoprotection.

## Methods

### Plasmids and LC3B mutant generation

The plasmid for mCherry-LC3B was kindly gifted by Dr. Sovan Sarkar (Massachusetts Institute of Technology) and GPR1/OA1-GFP construct was a kind gift from Dr. Elena Oancea from Brown University, USA. mCherry N1 and mCherry C1 were from Clontech Laboratories (632524, 632523).

SKIP-HA was a kind gift from Dr. Ivan Dikic and GST-SKIP (1-282) was generated by cloning the RUN domain containing 1-282 amino acids of SKIP by PCR amplification and cloning in pGEX-6P1 using EcoRI and SalI restriction sites. Mutant LC3B constructs in mCherry vector and SKIP-LIR domain deletion constructs in mammalian CMV-HA tag vector were generated using primers listed in **table S1**

Antisense oligonucleotides used to silence kinesins, LC3B or SKIP are listed in **table S1**

### Cell culture and RNAi screen

B16 mouse melanocytes were cultured in bicarbonate buffered DMEM-High Glucose (Sigma. D-5648), supplemented with 1x anti-anti (Gibco), 10% FBS (Gibco) at 37°c with 5% CO_2_. Cells were seeded in 24 well tissue culture plate and at 70% confluence transfected with a specific kinesin or non-targeting mouse si RNA (100nM) using Dharmafect II reagent (Thermo Scientific: T-2002-03 1.5 mL). si RNAs were custom synthesized against the chosen sense m RNA sequence of a specific mouse kinesin. All antisense oligonucleotides were synthesized by Thermo Fischer Scientific. Table 1 enlists all the si RNA sequences and their respective sense sequence. Gene complementation was performed by co-transfecting the cells with 2 µg of OA1-GFP. The cells were trypsinised and re-seeded after 24 hours of transfection for microscopic studies. The cells were imaged using Cellomics microscope, which involved acquisition of data from hundred fields per siRNA with 20X objective. The fields were then individually analysed and scored for perinuclear aggregation phenotype. This was repeated thrice, with 100 fields per siRNA, per experiment. The percentage of melanocytes showing perinuclear aggregation were categorised as +, ++ and +++ to indicate 20-40%, 40-60%, >60% of average cells showing melanosome clustering phenotype respectively.

Validation of knock-down phenotype observed upon silencing of 6 kinesins namely-– *Kif1a, Kif1b, Kif2b, Kif5b, Kifc1* and *Kifc3* was done by targeting these kinesins individually by SMARTpool siRNA designed against each of them. A high magnification confocal imaging was then done to affirm the role of these kinesins in melanosome transport.

### Immunocytochemistry

siRNA transfected cells were fixed for 10 minutes at 37°C in fresh fixation buffer consisting of 100 mM K-PIPES, 10 mM EGTA, 1 mM MgCl_2,_ 0.2% Triton X-100, 4% formaldehyde, pH to 6.9 with KOH. 0.1%. Triton X100 in TBS (TBS-TX) was used for washing. Blocking was done with 2% BSA in TBS-TX. Staining with HMB45 (DSS Imagetech (M0634)) and Tubulin (ab6046; 1:100) was done at room temperature for 2 hours in blocking buffer, rinsed 5 times with TBS-TX, incubated with secondary antibodies produced in mouse or rabbit (diluted 1:500 in blocking buffer) conjugated to Alexa fluor 488 and 568 for 1 hour at room temperature, and washed five times with TBS-TX. Nuclear staining was performed with SlowFade Gold Antifade mountant with DAPI (Invitrogen-S36938).

### Imaging

Fluorescent samples were visualised on a Zeiss LSM 510/ LSM 710 confocal system (built around a Zeiss Axiovert 200 M inverted microscope), using a 63X × 1.3 N.A. oil objective. Data were collected as z-stacks with approximately 25 planes and 0.5-0.6 µM spacing between each plane. The overlay quantification was performed on Zeiss co-localisation parameter by setting the background cut off values with single colour channels. The pixels that overlap over and above the background cut-off were considered as co-localized pixels. Individual overlap coefficient and Pearson’s coefficient was calculated for all pixels. Merged image was created using the maximum intensity projection software in-built in Zeiss system.

Secondary screen for the 6 shortlisted kinesin candidates was done using confocal imaging at 63X. The endogenous levels of each kinesin motor were individually disrupted in B16 cells using siRNA (as mentioned above) and stained with HMB45 and Tubulin to monitor perinuclear aggregation. Melanosome dispersion index (MDI) per cell was calculated as ratio between the raw integrated density of HMB45 staining in perinuclear region (region marked on basis of DAPI staining) by total raw integrated density of HMB45 in total cell (region marked based on tubulin staining).

### Melanosome preparation

B16 cells were grown to confluence and harvested using 0.1% trypsin-EDTA. The cells were then washed with 10% FBS and pelleted by centrifugation at 1000 g for 5 min at 4°C. Cells were injected in C57 black mice and the B16 melanoma was allowed to grow for 15 to 20 d. The tumor was excised and was washed in homogenization buffer (0.25 M sucrose [Merck, CAS 57–50–1], 10 mM HEPES, 1 mM EDTA, 2% antibiotic and antimycotic [ThermoFisher Scientific, 15240062], pH 7.2 and homogenized with 120 strokes of a Dounce glass homogenizer (Sigma, D8938). The homogenate was then centrifuged at 1000 g for 10 min at 4°C to prepare the post nuclear supernatant. The post nuclear supernatant was collected and separated on a stepwise sucrose density gradient and centrifuged at 100,000 g for 1 hour at 4°C in a swing-out rotor (Beckman coulter). The sucrose density gradients comprised 0.25, 0.8, 1.0, 1.2, 1.4, 1.6, 1.8, and 2.0 M sequentially layered with the highest density fraction on the bottom.

The stage III and IV melanosomes, which preferentially localized to the high-density sucrose fraction, were collected. The isolated fraction was diluted with melanosome wash buffer (0.25 M sucrose, 10 mM HEPES) and centrifuged at 12,000 g for 30 min at 4°C to separate the melanosomal pellet from the sucrose.

### GST Protein purification

GST-ΔN10 LC3B and GST-SKIP (1-282) plasmids were transformed into *E. coli* BL21DE3. A single colony was inoculated in 3ml LB broth and incubated overnight at 37°C, 200 RPM. Secondary cultures of 500ml were set-up using 1% inoculum from overnight grown culture and were incubated at 37°C, 200RPM until at OD of 0.6 was attained. 0.5mM Isopropyl β-d-1-thiogalactopyranoside (IPTG) was added to this culture for induction of protein synthesis and the culture were incubated at 18°C at 200RPM overnight. Cell pellet were collected by centrifugation at 11000 RPM for 5mins at 4°C. Pellet were washed with ice cold PBS and either stored at −80°C until further use or for further stems in pull down as explained above.

### Pull-down assay

Affinity purification was performed using GST-LC3B (Enzo Life Sciences) as the ligand. Melanosomal lysate was incubated with GST-LC3B and the interacting proteins were pulled down using glutathione-Sepharose 4B beads. The beads were washed with PBS. Equal volume of beads was loaded on the denaturing gel. Protein loading was normalised by Ponceau. The blots were probed with respective antibodies to check for interacting partners. KIF1B (Rb Kif1b #ab72108, dilution: 1:1000) and KIF5B (#ab15155, dilution: 1:1000). GST protein was used as a negative control in this experiment to assess non-specific interactions. Pull down with GST-SKIP (1-282) was also perform in similar manner. Anti-P62 antibody (ab56416; 1:1000) and anti-LC3B antibody (ab51520; 1:1000 for western blot) were used to assess pull-down.

### Ethics Statement

Fish experiments were performed in strict accordance with the recommendations and guidelines laid down by the CSIR Institute of Genomics and Integrative Biology, India. The protocol was approved by the Institutional Animal Ethics Committee (IAEC) of the CSIR Institute of Genomics and Integrative Biology, India (Proposal No 45a). All efforts were made to minimize animal suffering.

### Zebrafish Maintenance

Zebrafish used in this study were housed at the CSIR-Institute of Genomics and Integrative Biology following standard husbandry practices. Wildtype zebrafish larvae were obtained by pair wise mating of adult.

### Morpholino Injections

Morpholino (MO) oligonucleotide (Gene Tools, USA) were dissolved in nuclease free water (Ambion, USA) at a concentration of 1 mM according to the protocols recommended by Gene Tools. 1 mM stocks of MO oligos were stored at –80°C until further use. Working aliquots of MO oligos were prepared and stored at 4°C. Glass capillary (World Precision) micropipettes were pulled using Sutter Instrument (USA) and clipped appropriately to deliver 1–3 nl solution into 1–2 cell zebrafish larvae. Eppendorf Femtojet and Narshige micromanipulator were used for injection. Post microinjection, larvae were raised in E3 in 100 mm Petri dishes at a density 100 larvae per dish. The larvae were incubated at 28^0^C. Details regarding the morpholino sequences are provided in Table S1.

### Phenotype analysis

The Morphants were raised in dark conditions for the experiment till 3-4dpf and fixed using 4% Formalin. The larvae were then embedded in 2% methylcellulose for imaging. The Imaging was performed using Zeiss stemi 2000-C microscope fitted with axiocamICc1 camera. The images were then analyzed and segregated into dispersed, partial and aggregated categories.

### α-MSH induced dispersion

3-4 dpf morphant larvae were taken and kept in 200ul of E3 water in microfuge tube. α-MSH (1mM) was added to it in a final concentration of 15uM. The tubes were tapped and kept in dark in an incubator for 15 minutes. The embryos were then formalin fixed and subsequently analysed for dispersion defects.

### Live Imaging Setup and analysis

The live imaging was performed using Leica DMI 6000B inverted microscope. Dark conditioned morphants were used for the experiments. 40 mm tissue culture dishes (Orange scientific) were used, on this the larva was placed in a drop of E3 water and then the larva was overlaid with 2% Low melting agarose (LMA). The larva was placed with its dorsal side towards the 10X objective before the LMA solidified. The plate was then filled with E3 water and the humidifier was used when the temperature was set at 32^0^C so as the temperature of the E3 water was maintained at 26-28^0^ C. LAS V3.6 software was used to setup the live imaging in the brightfield mode, the time interval being 3 seconds for a total of 15 minutes. The experiment was then exported as TIFF images and video files were created in the quicktime mode.

The images at each minute interval were taken up for analysis using ImageJ. The initial image i.e., t0 was taken as the starting point and the percentage area of the melanosomes in the melanophores were measured. At least 3 melanophores from each embryo were measured.

### Co-Immunoprecipitation

SKIP-HA construct was used to facilitate precipitation of SKIP from B16 cell lysate using anti-HA antibody (H3663-100UL from sigma and dilution 1:500). B16 cells expressing HA-SKIP were lysed on ice, centrifuged for 5 mins at 4°C. 500µl lysate was diluted in 500 µl of lysis buffer and 5 µl of HA-agarose beads were added to the mix. The complex was then incubated on a rotator 10 RPM at 4°C for 1 hour. Beads were pelleted down by centrifuging for 2 mins at 750g at 4°C. Collected beads were then washed twice with lysis buffer and finally resuspended in 12 µl sample loading buffer. 4% of lysate volume taken in immunoprecipitation was loaded as input. Samples were then analyzed by western blot for the efficiency of LC3 co-immunoprecipitation. Cell lysates were prepared using NP40 lysis buffer (Invitrogen, ThermoFisher scientific, FNN0021) reconstituted with protease inhibitor cocktail.

### *In-vitro* LC3B Phosphorylation

Affinity purified GST-LC3B or GST-ΔN10 LC3B were incubated with kinase assay dilution buffer (ab139435) containing 10uM ATP, in 1:1 ratio for 10 mins at room temperature. To this mix 4 units of active PKA (Sigma Aldrich-P2645-400UN) was added and the reactions were incubated at 30*C for 90 minutes. To terminate the reactions the beads were washed with GST buffer containing 50 mM HEPES pH 7.4, 150 mM NaCl, 1 mM EDTA, 1 mM EGTA, 0.1% Triton-X-100, 10% Glycerol, 25 mM NAF, 10 µM ZnCl2 for 3-5 times at 2500g at 4°C.

### Site directed mutagenesis

The LC3B (phosphorylation site Thr12) in mCherry C1 and SKIP-LIR mutants in CMV-HA tag vector were generated using the agilent Quick Change II site directed mutagenesis kit (cat no. #200522). Mutant strand synthesis was carried out using Pfu Polymerase. The primers used for mutagenesis are listed in **table S1.**

### LC3B silencing and gene complementation

B16 cells cultured to a confluence of 70% in 6 well plate was co-transfected with siLC3B and various siRNA resistant rescue constructs as per the experimental requirement. 24hrs post transfection the cells were seeded on circular coverslips in 24 well plate. Cells then taken for immunocytochemistry as previously described for HMB45. Endogenous LC3B knockdown was assessed by western blot using anti LC3B antibody (ab51520; 1:1000) and complementation was assessed by anti-mCherry antibody (ab167453; 1:1000). HRP-Beta tubulin (ab21058, 1:10000) was used as for normalization.

### Dynamics of melanosome movements

B16 cells expressing OA1-GFP plasmid were cultured to a density of 50% confluence in µ-Slide 4 Well ibiTreat: #1.5 polymer coverslip (80426). 10uM alpha-MSH was used to stimulate the cells immediately before the video microscopy using Leica SP8 STED microscope at 63x, 1.4 NA oil immersion objective. Multiple fields containing single cells were marked and imaged in XYZT mode for 5 mins since stimulation. Time interval = 30 sec.

### œ-MSH mediated phosphorylation of LC3B

B16 cells were cultured at a density of 2 x 10^5^ cells/well of a 6 well plate. 1uM **œ-** MSH was added to the media to stimulate phosphorylation in a brief period of 30mins. Prior to this the cells were either pre-treated with H89 or not for 2 hours. After treatment with alpha-MSH cells were scraped on ice with NP40 buffer (Invitrogen) containing 1X PIC(Roche), 1X PhosphoSTOP (Roche), glycerophosphate (10mM), Sodium Pyrophosphate(10mM), Sodium Orthovandate (1mM) and PMSF (1mM). Following incubation on ice for 30mins cell lysates were centrifuged at 13000rpm for 30mins at 4°C. Supernatants were estimated for protein concentration and then store at −80°C until further use.

## Acknowledgements

This work was supported by the Council for Scientific and Industrial Research (CSIR), India, through grant (TOUCH-BSC0302) and (RegenX-MLP2008), Department of Biotechnology through the grant (GAP0182). YSJ acknowledges DBT for Research Fellowship. R.S.G. is a J.C. Bose Fellow of the Department of Science and Technology, Government of India (SB/S2/ JCB-038/2015). TNV acknowledges CSIR support through the Young Scientist Award (OLP1118). We thank Dr Ivan Dikic (Goethe University, Germany) for mentoring and providing SKIP constructs and scientific inputs for the biochemical aspect of the work.

## Supplementary figures

**Figure S1a:**
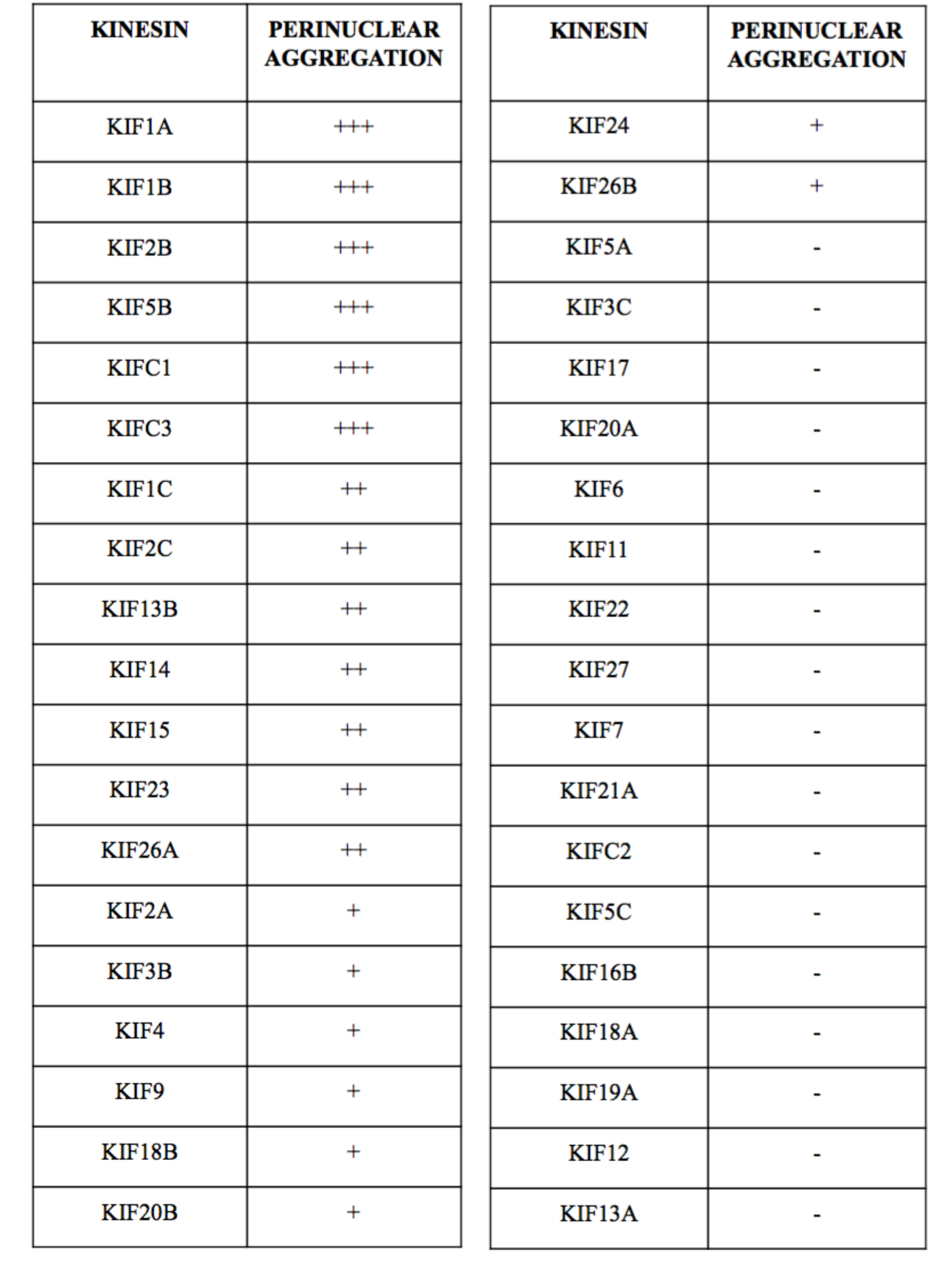
Silencing of individual kinesins lead to alteration in melanosome positioning in a non-redundant manner. 24 hours post transfection with respective siRNAs, B16 cells were fixed and labeled with fluorescent antibodies marking either the microtubule tracks (Tubulin-Alexfluor 488, Green) or melanosomes (HMB45-Alexafluor 561, Red). Images were captured using High content microscope by random sampling of 100 non-overlapping fields per siRNA at 20X magnification. Melanosome positioning as peri-centric or peripheral was analysed by bio-compartmetalization method. The ratio of HMB45 staining in the centre of the cell (Ci) to the intensity in the ring region (Ri) normalized to the respective area. List of siRNAs used in the screen is in supplementary table S 1.

**Figure S1B.**
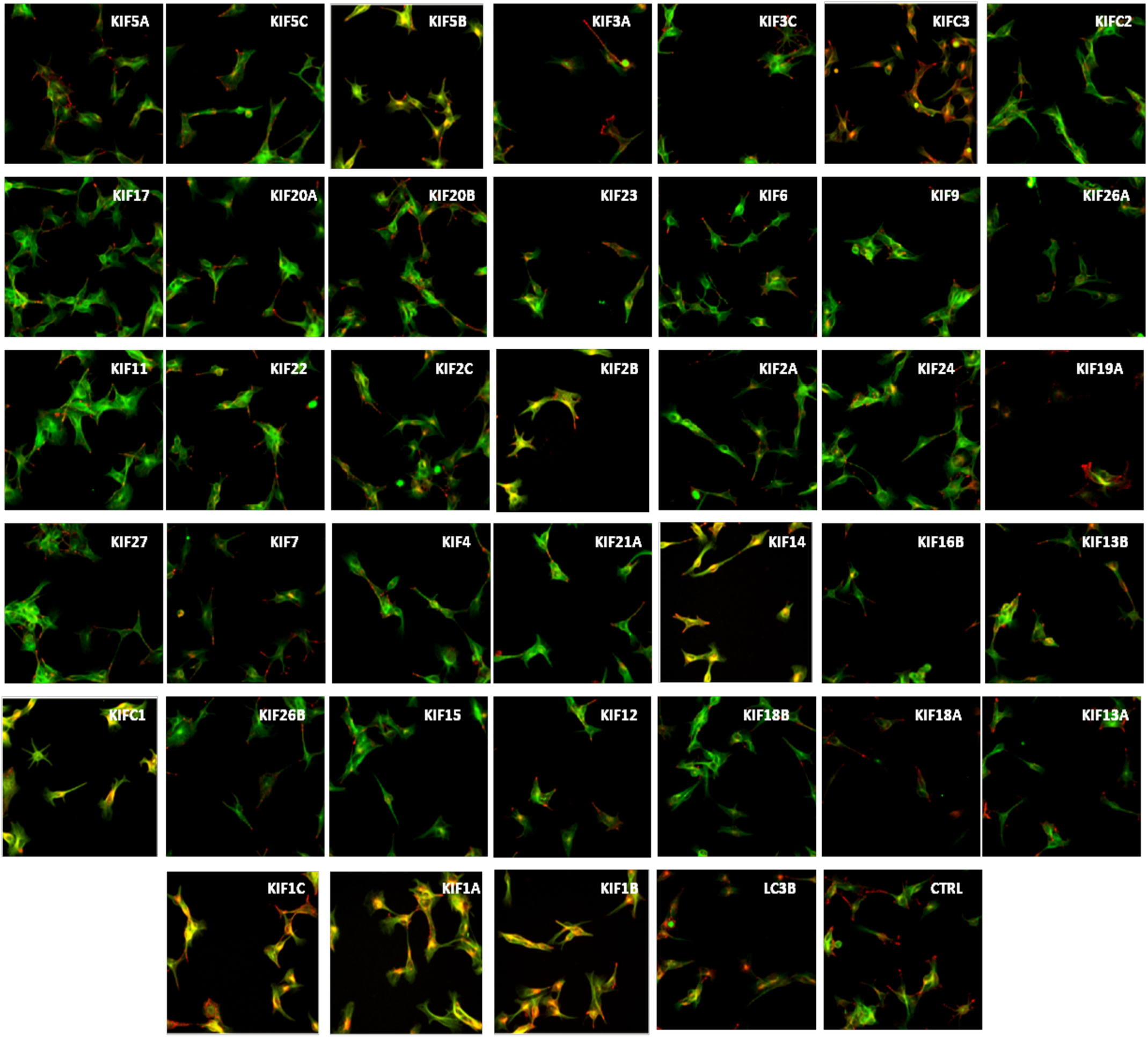
Summary of melanosome positioning effects of kinesin knockdown in B16. Melanosome distribution: +, ++ and +++ indicate 20-40%, 40-60%, >60% respectively, of the kinesin knocked down cells that exhibited perinuclear melanosome aggregation; n>100..

**Figure S2A:**
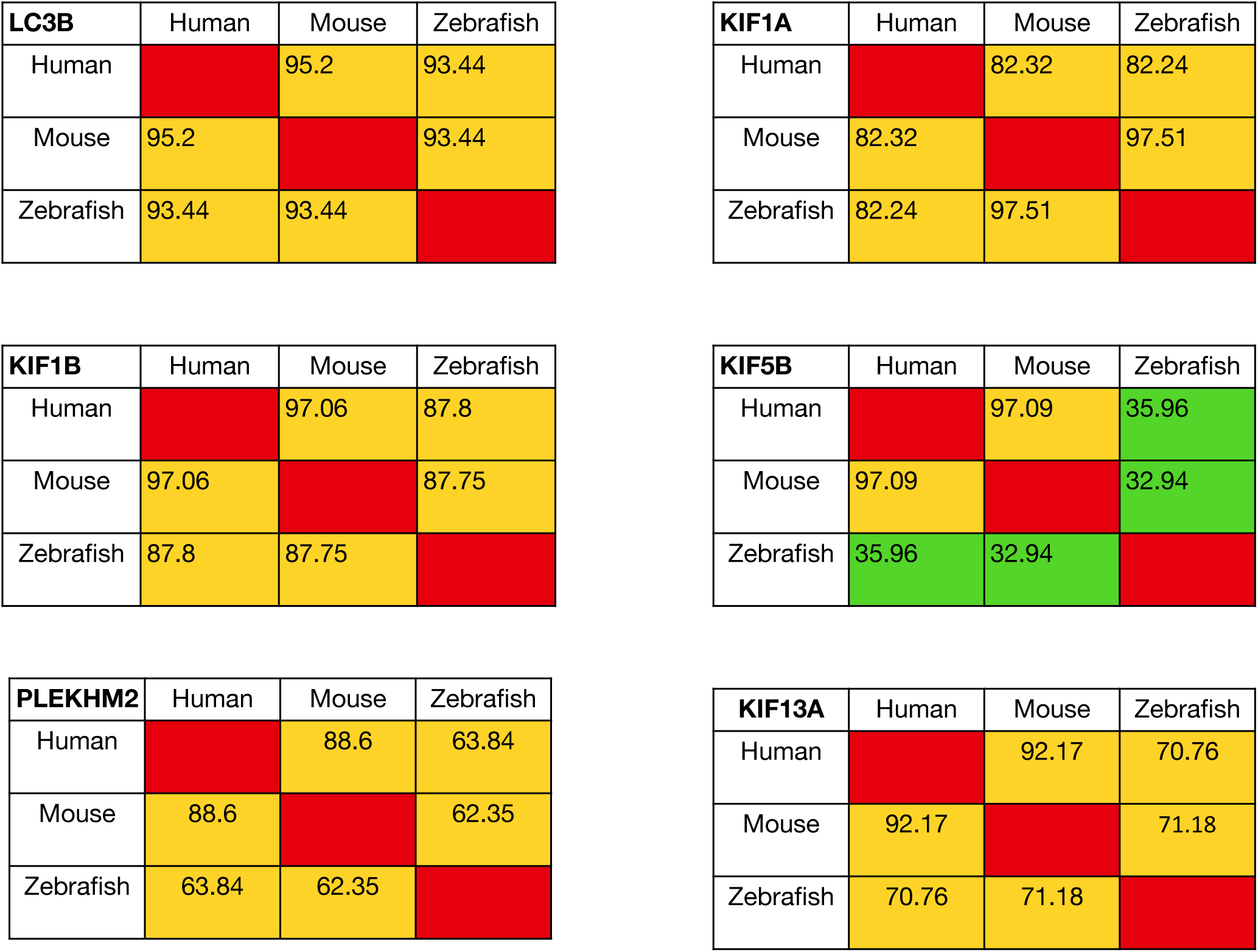
Protein involved in melanosome movement are conserved across species. Identity matrices for conservation of amino-acid sequences of kinesins-KIF1A, KIF1B, KIF13A and KIF5B; melanosomal tether-LC3B and the adaptor protein PLEKHM2 (SKIP)

**Figure S2B:**
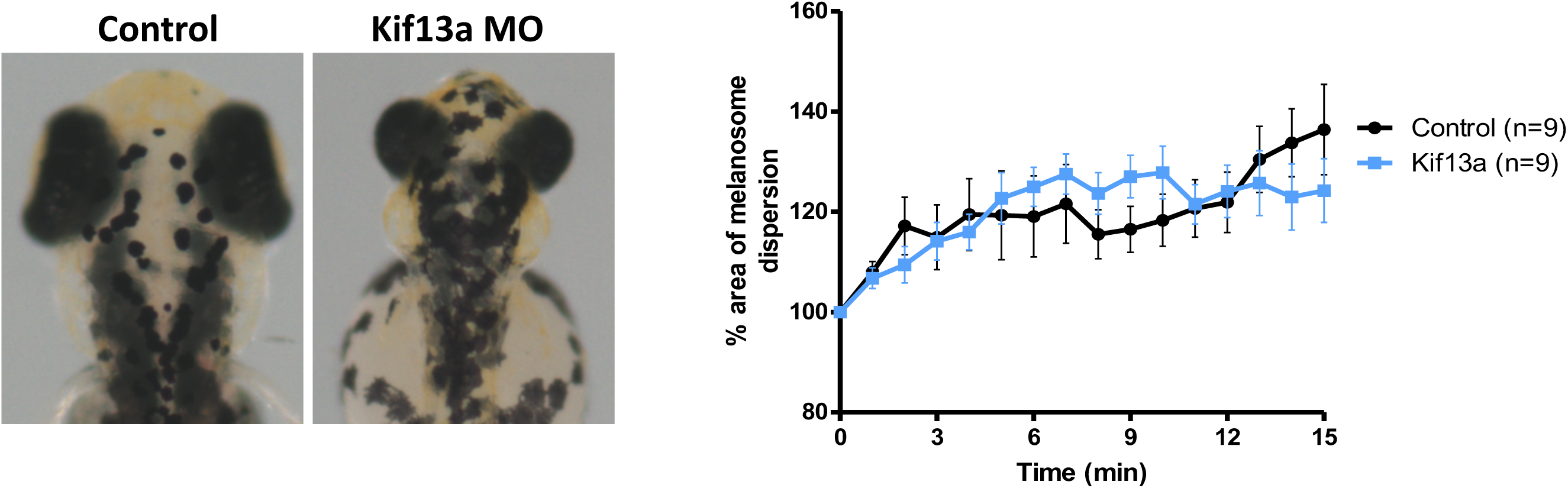
Silencing of Kif13a leads to reduced pigmentation without altering melanosome positioning in zebrafish embryos. Morpholino based silencing of kinesin involved in protein cargo transport to melanosomes, Kif13a resulted in no defects in melanosome spreading upon dark adaptation of injected embryos at 4 dpf. The graph depicts the live imaging-based melanosome dispersion analysis, control and kif13a morphants behave similarly, n=9 denotes the total number of melanophores analyzed per embryo. Control vs Kif13a MO = ns (pvalue=0.8427)

**Figure S3:**
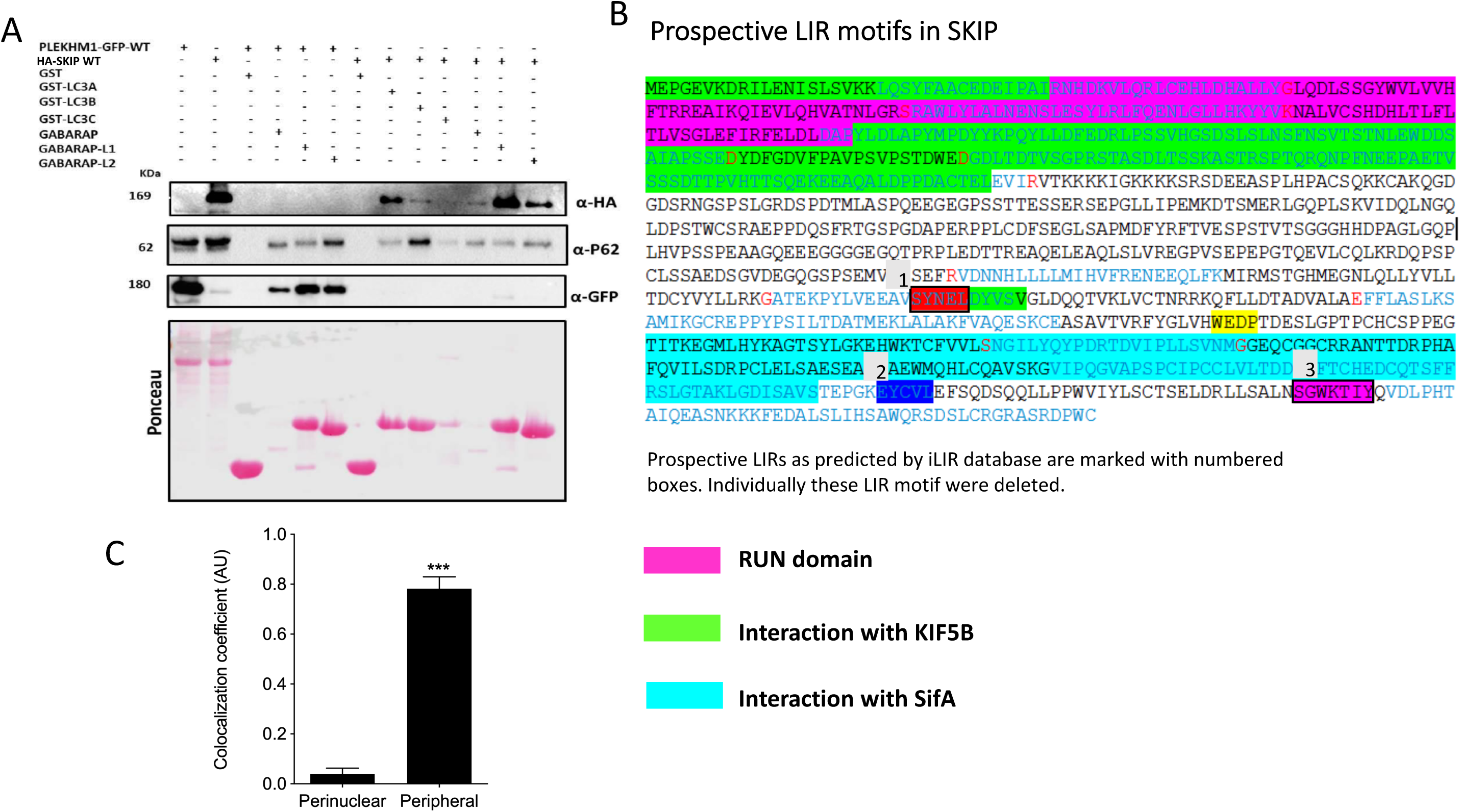
Association of adaptor protein with LC3 and related proteins. (A) Recombinant adaptor proteins PLEKHM1 (tagged with GFP) and SKIP (tagged with HA) were expressed in B16 cells and lysates were prepared for interaction study. Purified LC3A, LC3B, LC3C GABARAP L1 and GABARAP L2 were individually incubated with these lysates in combinations as indicated in the Figure. (B) Prospective LIR regions based on motif analysis from iLIR database on PLEKHM2 (SKIP) are indicated and were individually deleted to study their role in facilitating interaction between LC3B and SKIP. (C) Co-localization of SKIP(Alexafluor-488) with melanosome (HMB45-Alexafluor-561, pseudo-colored in purple) at cell center and cell periphery (Fig 4A) *** depicts p-value <0.0001 n=10 cells.

**Figure S4:**
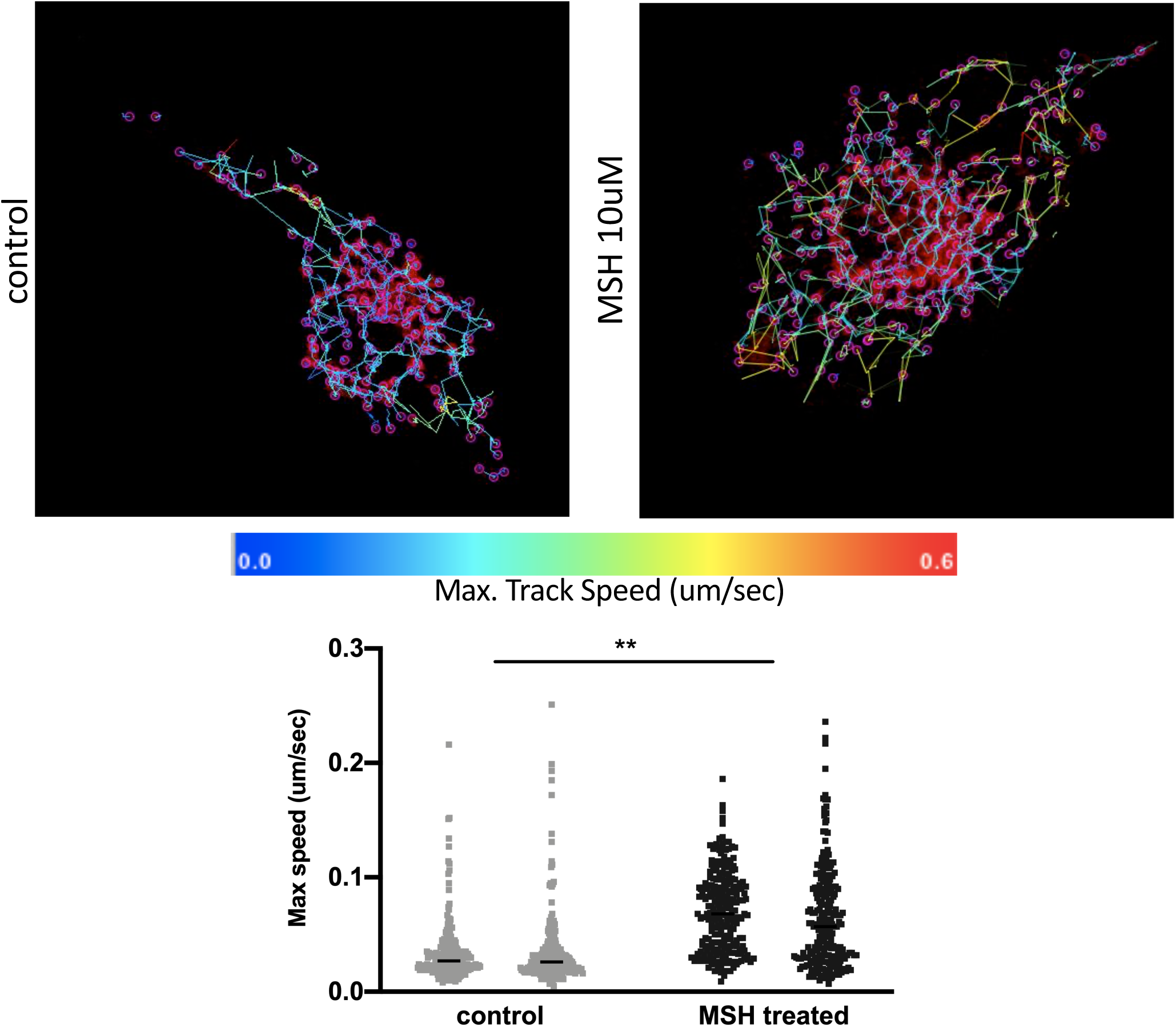
Speed of melanosome movement within cell increases upon treatment with α-MSH. B16 cells transfected with OA1-GFP (Melanosome marker, pseudo-coloured in Red) were imaged at 63x, 1.4 NA in XYCZT mode. Time interval between the images is 30 sec and total duration of imaging is 10mins. n= 2 cells per condition. Supplementary videos S5 and S6 are representative of melanosome tracking performed using Fiji plugin Trackmatev6.0.1. the track statistics generated by this analysis are given in supplement tables 3.1 to 3.4.

**Figure S5:**
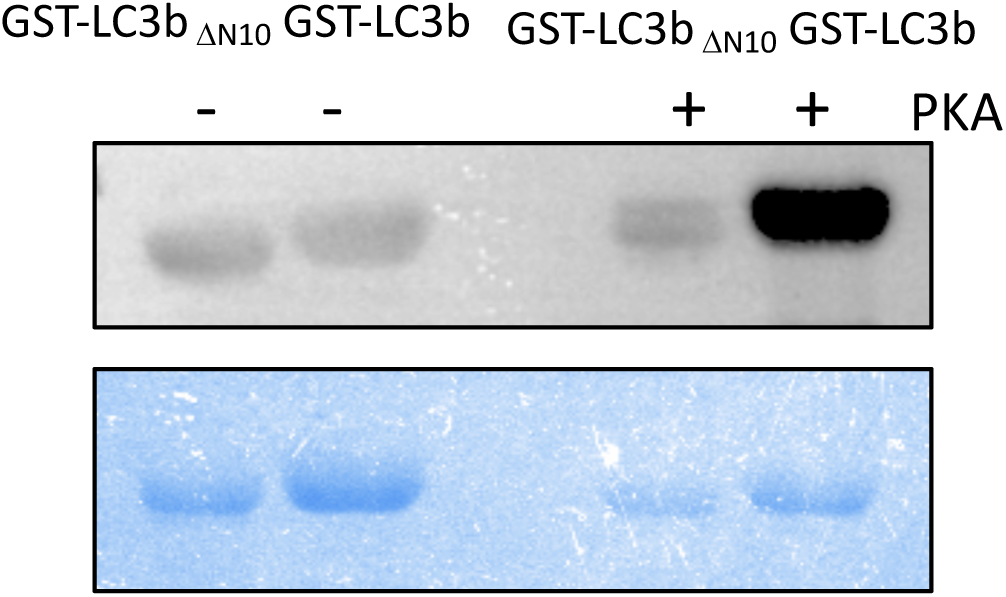
N-terminal amino acids are essential for PKA mediated phosphorylation on Thr12 of LC3B. Purified GST-LC3B, GST-DelN10-LC3B or control GST beads were treated with active PKA enzyme for 90mins at 30°C. Phosphorylation on Thr12 was then analysed by western transfer and probing of proteins with anti-phosphoThr12LC3 antibody (1:500).

